## Supplemental Information for "Navigating cross-reactivity and host species effects in a serological assay: A case study of the microscopic agglutination test for *Leptospira* serology"

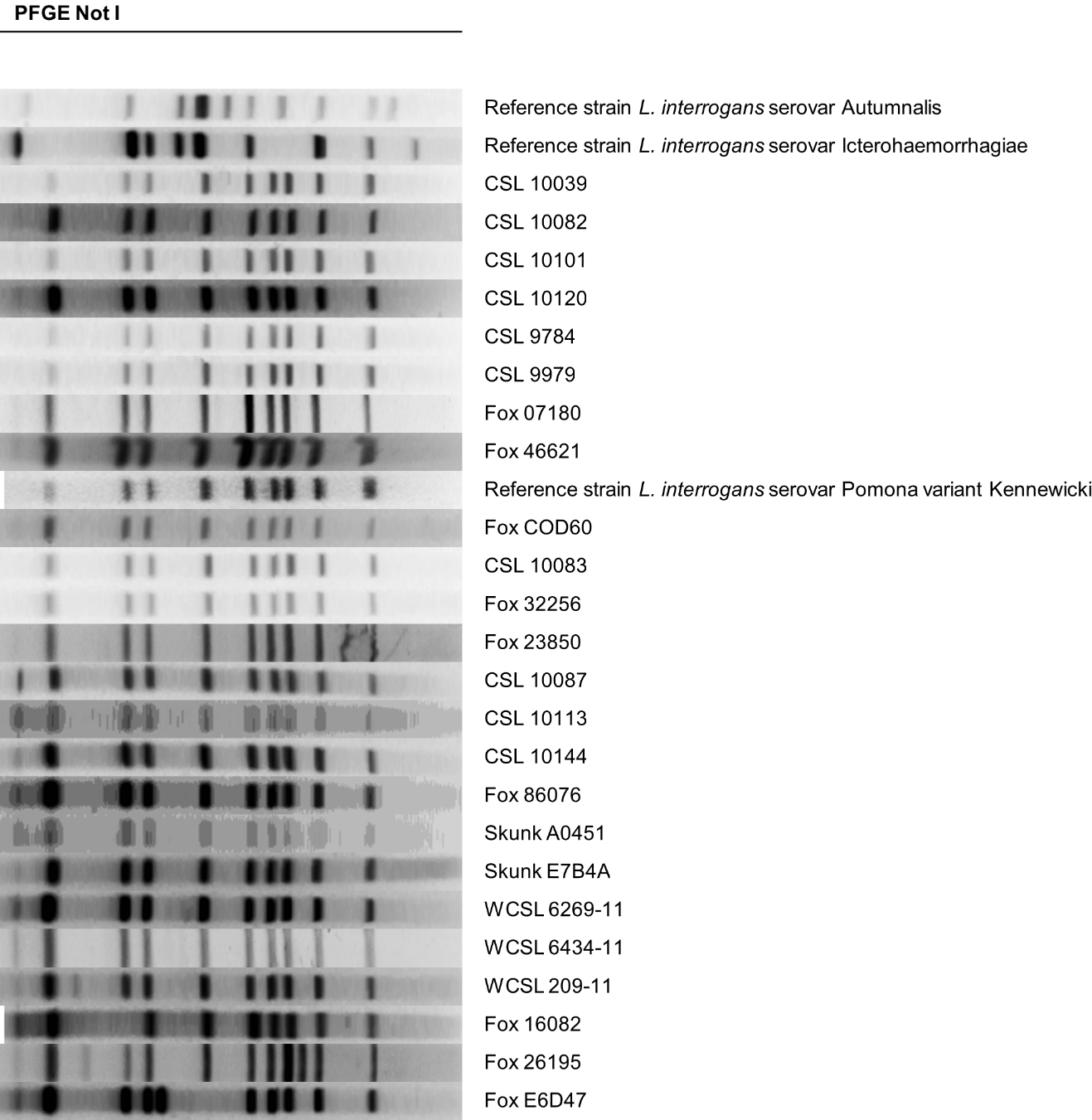

Figure S1. **Representative** **pulsed field gel electrophoresis (PFGE) results for 24 individual hosts (13 California sea lions, 9 island foxes, and 2 island skunks).** Labels beginning with CSL correspond to stranded sea lions, and WCSL corresponds to wild-captured sea lions. Three reference patterns for *L. interrogans* serovars Autumnalis, Icterohaemorrhagiae, and Pomona variant Kennewicki are also included. By convention, PFGE patterns that differ by three or fewer bands are considered the same serovar (Tenover et al., 1995); by this criterion, all samples tested in our study were classified as *L. interrogans* serovar Pomona variant Kennewicki.
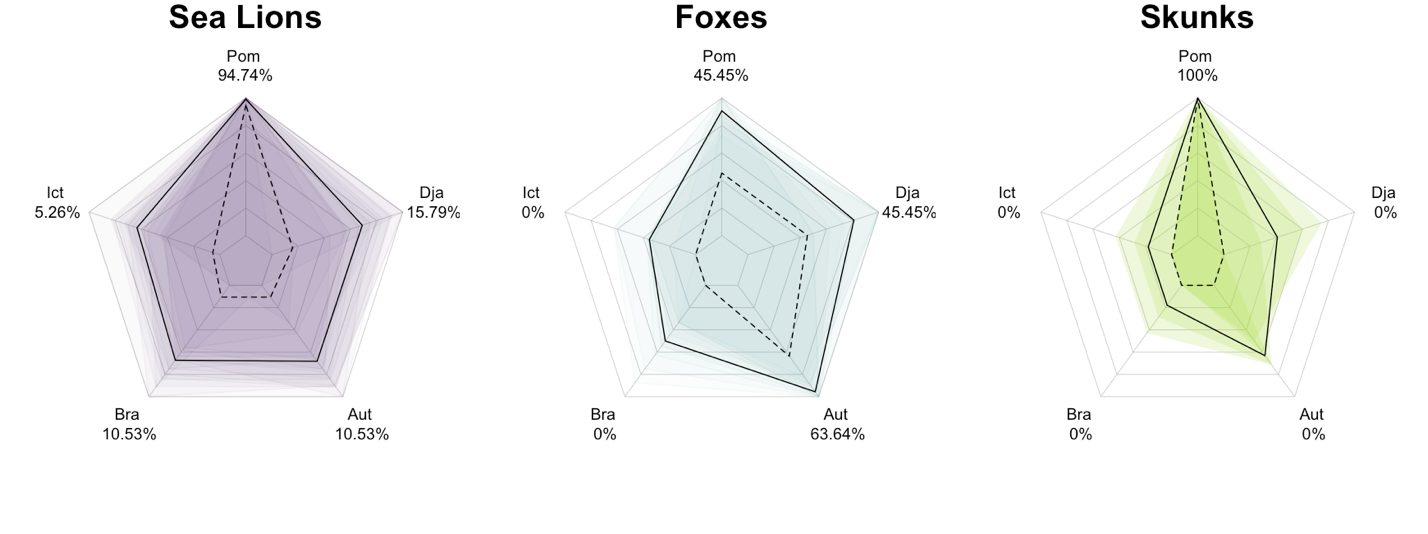

Figure S2. **Host-specific patterns of relative MAT antibody titers detected against five *Leptospira* serovars (Pomona, Djasiman, Autumnalis, Bratislava, and Icterohaemorrhagiae) when the infecting serovar is *L. interrogans* serovar Pomona, for individuals with positive PFGE only.** This is a replica of Figure 1 in the main text, except here only individuals with PFGE results are included. Each plot shows the relative antibody titer levels (antibody titer against one serovar divided by the highest antibody titer detected against any serovar in the 5-serovar MAT panel run for that sample) for California sea lions (left; purple; n=19), Channel Island foxes (middle; cyan; n=11), and spotted skunks (right; green; n=4). The shaded regions on each plot are a representative subsample of overlaid polygons, each linking the values for an individual sample. The continuous black line shows the relative antibody titer level for each sample (sample titer/maximum sample titer) averaged across all samples for each serovar for that species. The dashed black lines and the percentages associated with each serovar indicate the proportion of samples for which that serovar has the highest titer out of all serovars in that individual’s panel, regardless of the actual titer. These numbers add up to more than 100% for sea lions and foxes, since multiple serovars can have the highest titer for any given sample (e.g., a particular individual could have highest titer of 1:6400 against both Pomona and Icterohaemorrhagiae).

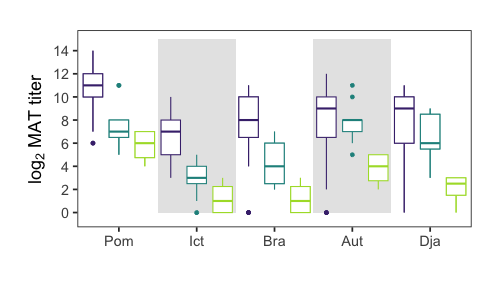

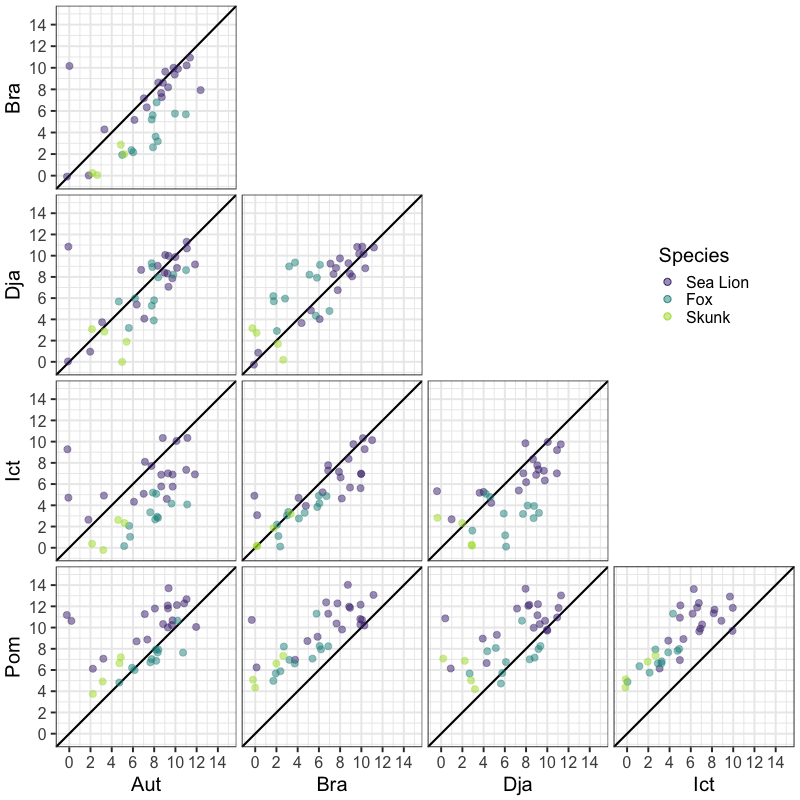

Figure S3. **Pairwise antibody titer levels against *Leptospira interrogans* serovars Pomona, Djasiman, Autumnalis, Bratislava, and Icterohaemorrhagiae in three host species, for individuals which are PFGE positive only.** MAT titers are shown as log_2_ dilutions. Statistical differences are shown in Tables S7-S10.

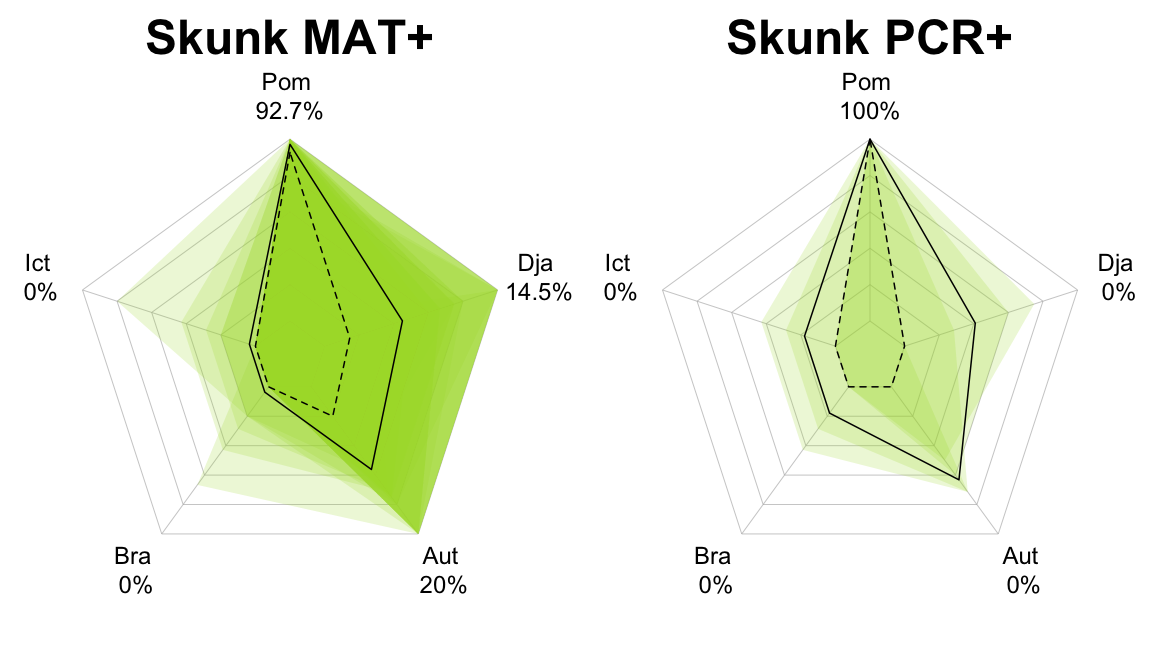

Figure S4. **Patterns of relative MAT antibody titers detected against five *Leptospira* serovars when the infecting serovar is *L. interrogans* serovar Pomona, for MAT-positive skunks (n=55) and PCR-positive skunks (n=4).**

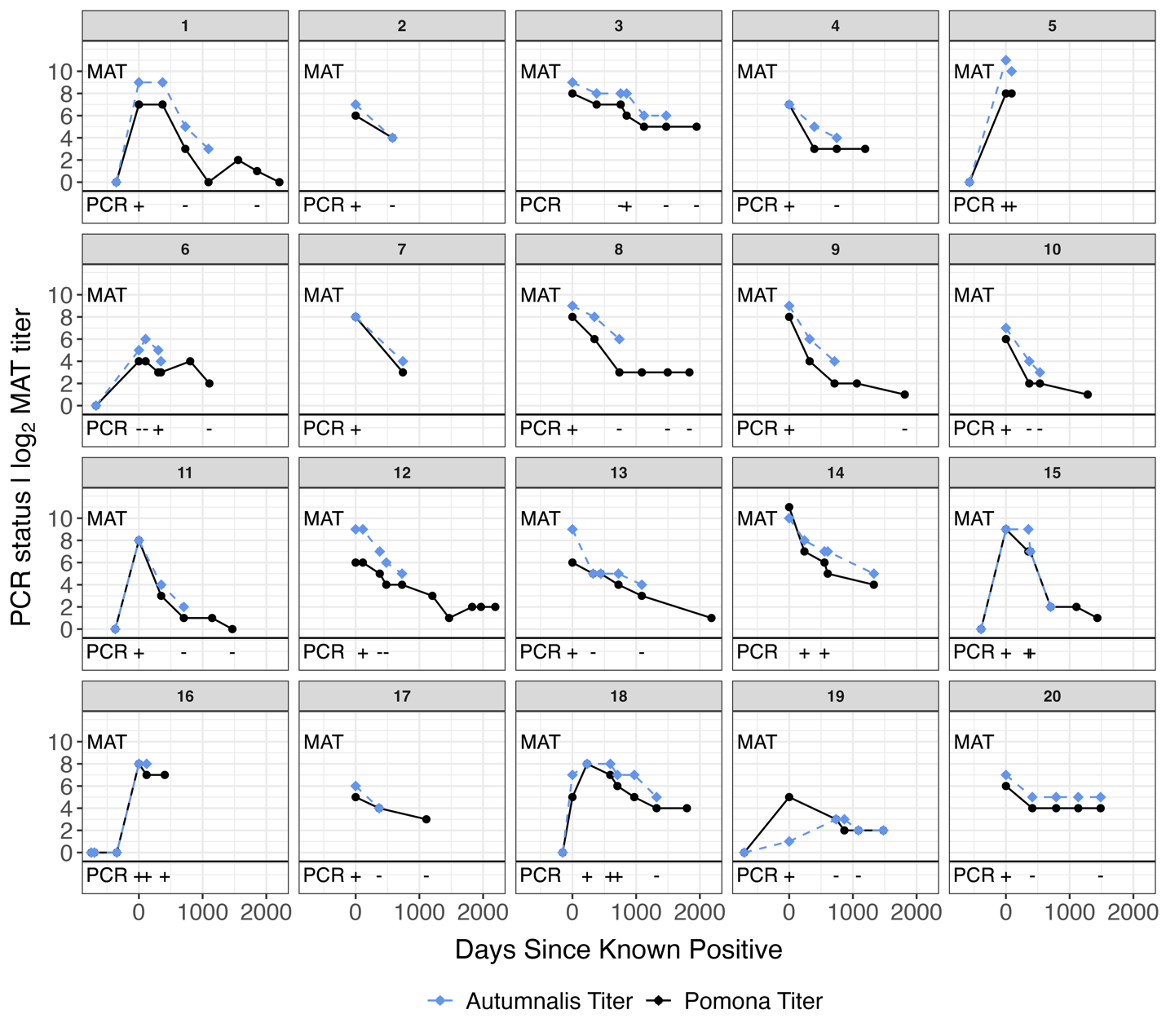

Figure S5. **Longitudinal antibody titer dynamics in Channel Island foxes.** Each facet illustrates the antibody dynamics of a single fox. The top panel of each facet shows antibody titers against *L. interrogans* serovars Pomona (black solid line) and Autumnalis (blue dashed line) from longitudinally collected serum samples. The bottom panel in each facet indicates the PCR test result from urine samples taken at the same time as serum collection.

**SUPPLEMENTAL TABLES**

Table S1. **The number of samples chosen per serovar Pomona titer level at Laboratory A for inter-laboratory titer comparison.**

| **Serovar Pomona titer level at Laboratory A** | 0 | 1:100 | 1:200 | 1:400 | 1:800 | 1:1600 | 1:3200 | 1:6400 | 1:12800 | 1:25600 | 1:51200 |
| --- | --- | --- | --- | --- | --- | --- | --- | --- | --- | --- | --- |
| **# Samples chosen for comparison** | 16* | 3 | 3 | 3 | 3 | 3 | 3 | 3 | 4 | 2 | 3 |

*6 samples were seronegative against Pomona but seropositive against Autumnalis; 10 samples were seronegative to both serovars

Table S2. **Pairwise ANOSIM statistics across host species.** Bolded values represent statistical significance.

| **Host species pairs** | **ANOSIM statistic** | **P-value** | **Adjusted p-value** |
| --- | --- | --- | --- |
| Fox vs. skunk | 0.786 | **0.001** | **0.003** |
| Fox vs. CSL | 0.257 | **0.001** | **0.003** |
| Skunk vs. CSL | 0.580 | **0.002** | **0.006** |

Table S3. **Kruskal-Wallis test statistics for differences among serovar within host species.** Bolded values represent statistical significance.

| **Species** | **Kruskal-Wallis chi-squared** | **df** | **P-value** |
| --- | --- | --- | --- |
| CSL | 152.23 | 4 | **< 2.2e-16** |
| Foxes | 166.41 | 4 | **< 2.2e-16** |
| Skunks | 11.73 | 4 | **0.019** |

Table S4. **Pairwise differences between serovar within a host species.** P-values are shown for each pairwise comparison. Bolded values represent statistical significance.

|  |  | *LogAut* | *LogBra* | *LogDja* | *LogIct* |
| --- | --- | --- | --- | --- | --- |
| CSL | *LogBra* | 0.171 | - | - | - |
|  | *LogDja* | 0.575 | 0.395 | - | - |
|  | *LogIct* | **3.6e-05** | **0.016** | **0.001** | - |
|  | *LogPom* | **4.6e-16** | **< 2e-16** | **< 2e-16** | **< 2e-16** |
| Fox | *LogBra* | **5.6e-15** | - | - | - |
|  | *LogDja* | **0.012** | **2.5e-11** | - | - |
|  | *LogIct* | **< 2e-16** | **1.8e-05** | **< 2e-16** | - |
|  | *LogPom* | **0.002** | **2.0e-12** | 0.571 | **< 2e-16** |
| Skunk | *LogBra* | 0.208 | - | - | - |
|  | *LogDja* | 0.330 | 0.601 | - | - |
|  | *LogIct* | 0.208 | 1.000 | 0.601 | - |
|  | *LogPom* | 0.301 | 0.095 | 0.095 | 0.095 |

Table S5: **Kruskal-Wallis test statistics for differences among host species for a given serovar.** Bolded values represent statistical significance.

| **Serovar** | **Kruskal-Wallis chi-squared** | **df** | **P-value** |
| --- | --- | --- | --- |
| LogPom | 70.44 | 2 | **5.05e-16** |
| LogDja | 8.07 | 2 | **0.018** |
| LogAut | 5.72 | 2 | 0.057 |
| LogIct | 81.77 | 2 | **< 2.2e-16** |
| LogBra | 36.87 | 2 | **9.87e-09** |

Table S6: **Pairwise differences between host species for a given serovar.** P-values are shown for each pairwise comparison. Bolded values represent statistical significance.

|  |  | *CSL* | *Fox* |
| --- | --- | --- | --- |
| LogPom | *Fox* | **1.3e-15** | - |
|  | *Skunk* | **0.0079** | 0.166 |
| LogDja | *Fox* | 0.227 | - |
|  | *Skunk* | **0.0305** | **0.005** |
| LogAut | *Fox* | 0.991 | - |
|  | *Skunk* | 0.062 | **0.006** |
| LogIct | *Fox* | **<2e-16** | - |
|  | *Skunk* | **0.003** | 0.125 |
| LogBra | *Fox* | **6.1e-08** | - |
|  | *Skunk* | **0.011** | **0.011** |

Table S7. **Kruskal-Wallis test statistics for differences among serovar within host species for PFGE positive individuals only.** Bolded values represent statistical significance.

| **Species** | **Kruskal-Wallis chi-squared** | **df** | **P-value** |
| --- | --- | --- | --- |
| CSL | 23.93 | 4 | **8.24e-05** |
| Foxes | 30.14 | 4 | **4.58e-06** |
| Skunks | 11.73 | 4 | **0.019** |

Table S8. **Pairwise differences between serovar within a host species for PFGE positive individuals only.** P-values are shown for each pairwise comparison. Bolded values represent statistical significance. Skunks are not shown in this table because they only have one PFGE-confirmed individual.

|  |  | *LogAut* | *LogBra* | *LogDja* | *LogIct* |
| --- | --- | --- | --- | --- | --- |
| CSL | *LogBra* | 0.853 | - | - | - |
|  | *LogDja* | 1.000 | 0.853 | - | - |
|  | *LogIct* | 0.230 | 0.230 | 0.230 | - |
|  | *LogPom* | **0.004** | **0.001** | **0.003** | **0.000** |
| Fox | *LogBra* | **0.002** | - | - | - |
|  | *LogDja* | 0.407 | **0.030** | - | - |
|  | *LogIct* | **0.000** | 0.310 | **0.002** | - |
|  | *LogPom* | 0.451 | **0.002** | 0.616 | **0.000** |

Table S9: **Kruskal-Wallis test statistics for differences among host species for a given serovar.** Bolded values represent statistical significance.

| **Serovar** | **Kruskal-Wallis chi-squared** | **df** | **P-value** |
| --- | --- | --- | --- |
| LogPom | 17.4 | 2 | **0.000** |
| LogDja | 9.34 | 2 | **0.009** |
| LogAut | 6.04 | 2 | **0.049** |
| LogIct | 21.31 | 2 | **2.36e-05** |
| LogBra | 14.02 | 2 | **0.001** |

Table S10. **Pairwise differences between host species for a given serovar.** P-values are shown for each pairwise comparison. Bolded values represent statistical significance.

|  |  | *CSL* | *Fox* |
| --- | --- | --- | --- |
| LogPom | *Fox* | **0.001** | - |
|  | *Skunk* | **0.007** | 0.107 |
| LogDja | *Fox* | 0.171 | - |
|  | *Skunk* | **0.017** | **0.017** |
| LogAut | *Fox* | 0.528 | - |
|  | *Skunk* | 0.082 | **0.016** |
| LogIct | *Fox* | **0.000** | - |
|  | *Skunk* | **0.004** | 0.082 |
| LogBra | *Fox* | **0.009** | - |
|  | *Skunk* | **0.015** | **0.034** |

Table S11. **Kruskal-Wallis test statistics across laboratories for serovars Pomona and Autumnalis.** Bolded values represent statistical significance.

| **Serovar** | **Kruskal-Wallis chi-squared** | **df** | **P-value** |
| --- | --- | --- | --- |
| LogPom | 15.93 | 2 | **0.000** |
| LogAut | 2.87 | 2 | 0.238 |

Table S12. **Pairwise differences between laboratories for serovar Pomona.** P-values are shown for each pairwise comparison. Bolded values represent statistical significance.

|  |  | *Laboratory A* | *Laboratory B* |
| --- | --- | --- | --- |
| LogPom | *Laboratory B* | **0.028** | - |
|  | *Laboratory C* | **0.001** | **0.024** |
